# Response to nitrogen addition reveals metabolic and ecological strategies of soil bacteria

**DOI:** 10.1101/107961

**Authors:** Md Sainur Samad, Charlotte Johns, Karl G. Richards, Gary J. Lanigan, Cecile A. M. de Klein, Timothy J. Clough, Sergio E. Morales

## Abstract

The nitrogen (N) cycle represents one of the most well studied systems yet the taxonomic diversity of the organisms that contribute to it is mostly unknown, or linked to poorly characterized microbial groups. While progress has allowed functional groups to be refined, they still rely on *a priori* knowledge of enzymes involved, and the assumption of functional conservation, with little connection to the role the transformation plays for specific organisms. Here, we use soil microcosms to test the impact of N deposition on prokaryotic communities. By combining chemical, genomic and transcriptomic analysis we are able to identify and link changes in community structure to specific organisms catalyzing given chemical reactions. Urea deposition led to a decrease in prokaryotic richness, and a shift in community composition. This was driven by replacement of stable native populations, which utilize energy from N-linked redox reactions for physiological maintenance, with fast responding populations that use this energy for growth. This model can be used to predict response to N disturbances and allows us to identify putative life strategies of different functional, and taxonomic, groups thus providing insights into how they persist in ecosystems by niche differentiation.

## Introduction

Modern microbiology techniques have given us unprecedented access to the microbial world (Spiro 2012; Rinke *et al*. 2013), yet soil microbial communities remain poorly understood (Delmont *et al*. 2015). While many studies have focused on the diversity or abundance of key populations (Taylor *et al*. 2012; Wei *et al*. 2015; Gubry-Rangin *et al*. 2015), fewer have looked at the transcriptional profiles over time (Nicol *et al*. 2008; Morales & Holben 2013), and even less have done so for multiple groups at the same time (Liu *et al*. 2010; 2014; Brenzinger *et al*. 2015). This is particularly true of organisms involved in nitrogen (N) cycling in soils. The complexity of the underlying processes combined with the diversity of microbes contributing to each process provides a large challenge to identifying mechanisms active at any given time (Butterbach-Bahl *et al*. 2013). Currently we lack enough information to understand basic ecological concepts linked to N cycling *in situ* such as: i) substrate competition at both inter and intra species level, ii) full diversity of both present and active N cycling populations, iii) and the life strategies of these populations which in turn control their responses (both as observed growth or transcriptional changes).

The initial discovery of ammonia oxidizing archaea (AOA) and recognition as important players in the N cycle (Leininger *et al*. 2006; Hatzenpichler 2012; Stahl & la Torre 2012) highlighted the unexpected gaps in knowledge. Later studies have suggested different life strategies for AOA when compared to ammonia oxidizing bacteria (AOB) (Sterngren *et al*. 2015), but this may be complicated by variance across strains (Bayer *et al*. 2015). One major unknown is whether observations made in studies, or organisms, from one ecosystem translate to others.

It is well established that individual intermediates in the N cycle can be used for specific reasons (i.e. ammonia oxidation provides electrons, while denitrification intermediates accept reducing equivalents), but the purpose of the reactions for any organism is another major unknown. That is, while some organisms carry out these processes for electrogenic purposes that can result in growth, others do it in order to maintain redox homeostasis (e.g. to dissipate excess reductants) (Green & Paget 2004). Unfortunately examples where an organism harbours multiple versions of the same enzyme for completely different purposes (respiration vs. redox balance) exist (Hartsock & Shapleigh 2011), and are likely to limit generalizations.

Despite this, studies focusing on population changes in response to manipulations have consistently recorded conserved patterns (e.g. growth of AOB but not AOA (Jia & Conrad 2009; Di *et al*. 2009; Pratscher *et al*. 2011)) suggesting that responses by specific populations in a given location or ecosystem are predictable. However, the debate continues on whether niche specialization and differentiation can be determined based solely on correlations, without analyzing the wider array of processes that contribute or influence any given N transformation (Prosser & Nicol 2012). This is relevant in ecosystems such as agricultural grassland where an understanding of N cycling is crucial for management of both productivity and greenhouse gases (Herrero *et al*. 2016), of which nitrous oxide (N_2_O) is a key player (Reay *et al*., 2012).

In grazed pastures (i.e. agricultural grasslands) N deposition through ruminant urine drives the emissions of N_2_O (Saggar *et al*. 2013). In this system a full cascade of transformations begin with urea and can result in accumulation of any intermediate depending on conditions, but with a final end product of N2 or N_2_O. While the chemical transformations have been explored (Hamonts *et al*. 2013; Baral *et al*. 2014; de Klein *et al*. 2014a; b), mechanistic understanding of the populations catalyzing the reactions, and the purpose they serve for the organisms is less clear. In this study, we aimed to identify active N-transformation pathways as well as changes in microbial populations/taxa abundance and transcriptional activity for organisms involved in N loss (through gases) in response to urea (simulated ruminant urine deposition event) and varying moisture content. Observed chemical transformations were linked to changes in genotype (functional potential through DNA; a proxy for population changes), expression of genotype (RNA profiles), and total community composition (specific taxonomically defined populations based on the 16S ribosomal rRNA gene). We hypothesized that sequential transformation of nitrogenous intermediates would be coupled to changes in expression of functional genes catalyzing production and consumption of intermediates. Alternatively, transformations not linked to population, or expression changes, would be driven by other (abiotic) pathways. We also hypothesized that changes in transcription, or population size, could serve to determine life strategies of microbes utilizing each intermediate (whether they are used for growth vs. physiological maintenance). To test this we mimicked a ruminant urine-N deposition event using repacked soil cores (soil bulk density= 1.1 Mg m-3) on tension tables monitored for 63 days. Soils were treated with urea under two different moisture contents: high (near saturation; -1.0 kPa) and low (field capacity; -10 kPa) moisture. Simultaneous measurements of soil chemistry, gas kinetics, microbial community composition (by 16S rRNA gene amplicon sequencing) and functional gene abundance (for nitrification and denitrification) at DNA (gene) and RNA (transcript) levels were performed to determine the active populations and pathways.

## Materials and methods

### Sample collection and experimental design

A detailed methodology can be found in (Clough *et. al*., In review). In brief, soil was collected from a permanently grazed agricultural grassland (dairy pasture) in March (early spring) at the Teagasc Moorepark Research Centre, County Cork, Ireland (8°15’W, 52°9’N). The soil is classified as a Typical Brown earth from the Clashmore Series (Gardiner & Radford 1980). Soil was sampled after the turf was removed and a spade was used to randomly sample the A-horizon (5-20 cm depth, excluding grass layer). To avoid fresh N loading, fields had not been grazed for over a month. Field moist samples were immediately shipped to Lincoln University, New Zealand and kept at 4oC until processed. Prior to use, soil was sieved (≤ 2 mm) to remove any stones, plant roots or earthworms and packed into stainless steel rings (7.3 cm internal diameter, 7.4 cm deep) to a depth of 4.1 cm at *in situ* soil bulk density (1.1 Mg m^-3^ with a gravimetric water content (*θg*) of 0.24 g water g^-1^ soil). The resulting cores had a total porosity of 0.58 cm^3^ pores cm^-3^ soil and were arranged in a factorial experiment replicated four times. Soil cores were maintained at two moisture contents: high (near saturated; -1.0 kPa) and low (field capacity; -10 kPa) moisture using tension tables (Romano et al., 2002). These moisture contents, -1 and -10 kPa respectively, corresponded to 53% and 30% volumetric water content, or 91% and 52% water-filled pore space (WFPS). Nitrogen was applied as a urea solution at 2141 kg urea/ha dry soil (equivalent to a single urination event at the higher rate expected under bovine urine deposition of 1000 kg N ha^-1^). Four treatments in total were carried out (replicated four times each for a total of 112 cores analyzed) representing two levels of urea and two levels of moisture: urea + high moisture (HM +N; Urea _- 1.0kPa), urea + low moisture (LM +N; Urea _-10kPa), no urea + high moisture (HM–N; No Urea _-1.0kPa) and no urea + low moisture (LM –N; No Urea_-10kPa). All cores where held at 20oC for a period of 63 days.

### Soil pH, and inorganic-N measurements

Soil pH was monitored throughout the experiment using a flat surface pH electrode (Broadley James Corp., Irvine, California). Inorganic N concentrations (NH_4_^+^, NO_2_^-^, NO_3_^-^) were determined by destructively sampling batches of soil cores. Each core was homogenized and a subsample was extracted (10 g dry soil: 100 ml 2M KCl shaken for 1 hour), filtered (Whatman 42) and analyzed using flow injection analysis (Blakemore et al., 1987). N_2_O flux was determined by placing a soil core into a 1-L stainless steel tin fitted with a gas-tight lid and rubber septa. The headspace was sampled after 15 and 30 minutes and analyzed using an automated gas chromatograph (8610; SRI Instruments, Torrance, CA), linked to an autosampler (Gilson 222XL; Gilson, Middleton, WI) as previously described (Clough et al., 2006).

### Nucleic acids extraction

Samples for RNA and DNA extraction were collected simultaneously with samples for inorganic N analysis, but only samples at 0, 7, 14, 21, 35, 63 days were processed for nucleic acids. Each biological replicate was extracted and analyzed separately. For each extraction 2 g (wet weight) of soil were processed using the PowerSoil Total RNA Isolation and DNA Elution Accessory Kits (MoBio, Carlsbad, CA) as per manufacturer’s instructions, with slight modifications. Bead beating was done in a Geno/Grinder 2010 (SPEX SamplePrep, LLC, Metuchen, NJ) using two rounds of beating (1750 strokes/min) for 15 s with a 1 min pause in between. The total elution volume for RNA and DNA was 60 µl and 100 µl respectively. RNA was treated with DNase I (RNase-Free) (New England Biolabs, USA) as per the manufacturer’s protocol. RNA quality was assessed by denaturing gel electrophoresis. RNA and DNA concentration, purity and humic acid contamination were determined using a Nanodrop Spectrophotometer, ND-1000 (Thermo Scientific). All extractions were stored at -80 °C until downstream analyses.

### Reverse transcription (RT)

Triplicate cDNA conversions (technical replicates) were performed for each RNA extraction using the Maxima H Minus First Strand cDNA Synthesis Kit (Thermo Scientific) according to manufacturer’s protocol. Each 20 µl reaction contained: 13 µl of RNA (208 ng Total RNA), 1 µl of random hexamers (100 pmol), 1µl of dNTP mix (0.5 mM final conc.) and 5 µl of master mix (4 µl of 5X RT buffer and 1 µl Maxima H Minus reverse transcriptase). All technical replicates for a sample were combined and stored at -80°C until further analysis. All further analyses were performed on the same cDNA pool for each sample.

### 16S rRNA gene amplicon sequencing

16S rRNA gene amplicon sequencing was performed using primers 515F/806R (V4 region of the 16S gene) and the Earth Microbiome Project conditions (Version 4_13) (Caporaso *et al*. 2012). All samples were run simultaneously on a single Illumina MiSeq run. Sequences were first processed in Qiime (version 1.9.1) using default parameters (Caporaso *et al*. 2010). Sequences were clustered into Operational Taxonomic Units (OTUs) at 97% sequence similarity using the SILVA (version 119) reference library (Quast *et al*. 2012) and UCLUST (Edgar 2010) following the open-reference Operational Taxonomic Unit (OTU) picking protocol. Taxonomic identification was done using BLAST against the SILVA database (max-e value = 0.001) (Altschul *et al*. 1990). Subsampling and rarefactions (10 times) were performed to equal read depths of 7,400 per sample, and samples below that threshold were removed. After rarefaction, all 10 OTU tables were merged and exported for further processing in R (R Development Core Team 2008). The 16S amplicon sequences are available in the NCBI SRA database (accession numbers SRP091980).

### Quantification of gene and transcript abundance

Quantitative PCR (qPCR) was performed in 384-well plates using the ViiA7 real-time PCR system (Applied Biosystems, Carlsbad, CA). Absolute quantification was done using a 10-fold serial dilution (10^8^ to 10^1^) of known copy numbers of pGEM-T easy (Promega, Madison, Wisconsin, USA) cloned templates as standards. For all targets qPCR runs included cloned standards, no template control and no reverse transcription controls (RNA) run in triplicate. No inhibition or positive amplification on negative controls was observed for any target. All DNA and cDNA samples were run in quadruplicates to determine abundance of: prokaryotes (16S rRNA gene), ammonia oxidizers (archaeal [AOA] & bacterial [AOB] ammonia monooxygenase gene; *amoA*), denitrifiers (cytochrome cd1-type nitrite reductase gene; *nirS*, and Clade I nitrous oxide reductase gene; *nosZI*) and nitrogen fixers (nitrogenase gene; *nifH*).

All reactions were performed in 10 µl volumes containing: 1× Master Mix (Fast SYBR Green Master Mix, ABI), 0.2-0.6 µM of each primer [0.2 µM for AOA (Tourna *et al*. 2008), 0.6 µM for AOB (Rotthauwe *et al*. 1997; Avrahami *et al*. 2003); 0.5 µM for 16S rRNA (Hartman *et al*. 2009); *nirS* (Throbäck *et al*. 2004; Yergeau *et al*. 2007), *nosZI* (Henry *et al*. 2006) & *nifH* (Rösch & Bothe 2005)], 2 µl of template [DNA (1 ng total) or cDNA (80× diluted RT reaction, i.e. total 0.13 ng RNA)] and autoclaved Milli-Q H_2_O to a final volume of 10 µl. Primers and qPCR conditions are summarized in Table S1. A melt curve analysis (95°C for 15 s, 60°C for 1 min then increasing 0.05°C/s (data acquisition) until 95°C) was performed to test for specificity and to confirm no amplification in the negative controls.

### Statistical analyses

All statistical analyses were performed in R (R Development Core Team 2008) using the phyloseq (McMurdie & Holmes 2013), pvclust (Suzuki & Shimodaira 2006), vegan (Oksanen *et al*.) and mpmcorrelogram packages. Detailed descriptions can be found in supplemental methods.

### Growth rate estimation and prediction of rRNA operon (rrn) copy numbers

*rrn* copy numbers for identified OTUs were predicted using the ribosomal RNA operon copy number database (rrnDB) (Stoddard *et al*. 2015). For each OTU, information from the closest strain available was selected. In instances where a closely related organism was not available, the mean copy number for the closest taxonomic group (i.e. genus, class, etc.) was used. Copy numbers where then compared to the maximum observed abundance and the maximum observed fold change (calculated based on lowest observed abundance for the same organism in a preceding time point for OTUs showing growth or succeeding time points for those decreasing in abundance). An estimated growth rate was calculated for OTUs showing increases in population size in response to N using the following formula:

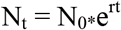

where: N_t_: The amount at time t; N_0_: The amount at time 0; r: exponential growth rate;

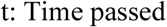

### Fit model for rrn copy numbers

Both non-linear (Michaelis-Menten) and linear regressions were used to fit *rrn* copy numbers and population changes (i.e. maximum abundance and fold-change), and growth rate (per day). The fit model was performed in R using “drc” and “ggplot2” packages.

## Results

### Soil pH and N transformation dynamics in response to urea

Soil pH increased from acidic (pH = 5.5 ± 0.1, i.e. mean ± SD) to alkaline reaching a maximum (pH = 8.7 ± 0.2) at day 3 in urea treated soils. Return to baseline pH was modulated by soil moisture with high moisture (HM; -1.0kPa) soil reaching baseline at day 35 and low moisture soils (LM; -10kPa) doing so at day 53 (Fig. 1). This shift in pH was linked to a successive N transformation process initiated with urea hydrolysis and leading to nitrification and denitrification: urea 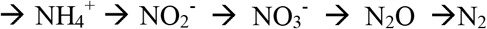 (Fig. 1). Sequential peak activity was observed for each transformation with the response modified by moisture. Maximum production (mean µg N g^-1^ soil) for each transformation was observed at day 3, 21 and 35 respectively for NH_4_^+^ (HM+N = 1758; LM+N= 1730), NO_2_^-^ (HM+N = 79.2; LM+N= 39.7) and NO_3_^-^ (HM+N = 429.2; LM+N= 335). Two distinct production peaks were observed for N_2_O, with a short pulse (0 to 5 days) reaching a maximum at day 2 for HM soils (11602.8 µg m^-2^ h^-1^) and day 3 for LM soils (46.8 µg m^-2^ h^-1^) (Fig. 1 and Supplementary Fig. S1). A second, longer duration (10 to ∼50 days), N_2_O pulse reached a maximum at day 28 for HM soils (6405.1 µg m^-2^ h^-1^) and day 30 for LM soils (448.9 µg m^-2^ h^-1^). The large N_2_O spike (first peak) between days 0 to 5 in the HM+N treatment was about 11.6% of the total N_2_O cumulative flux over 63 days, whereas in the LM+N treatment the 0 to 5 day periods accounted for 22.3% of the total N_2_O cumulative flux over 63 days.

**Fig. 1.**
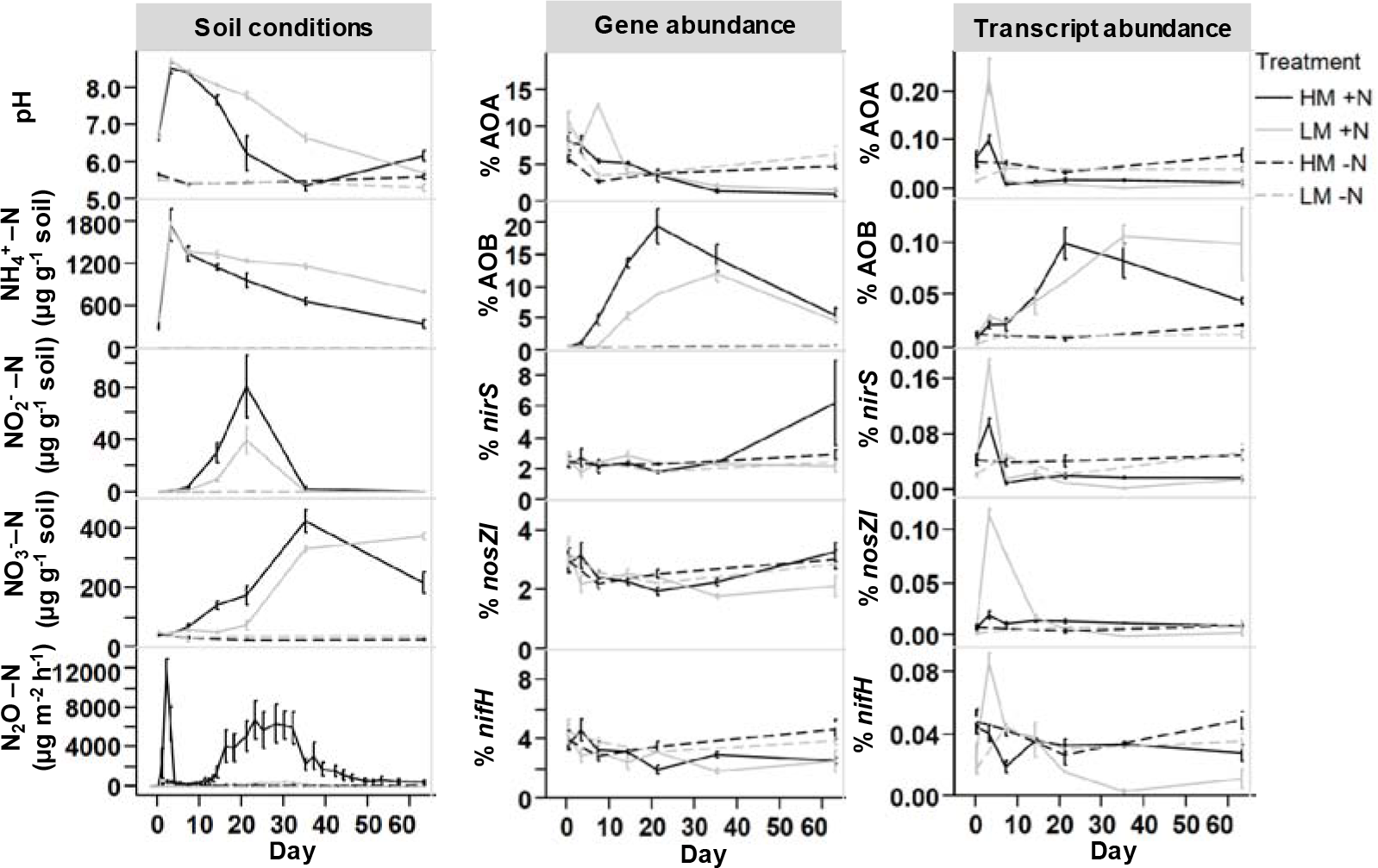
Chemical transformations and biological (functional group) response in soils treated with urea (+/- 1000 µg N/g dry soil) under two moisture conditions (LM = low moisture [-10kPa]; HM = high moisture [-1.0kPa]). Error bars are the standard error of the mean (n ≥ 3, except gene abundance data of day 7 [n=1; LM soil] and day 21 [n=1; LM soil]) for replicate mesocosms. Gene and transcript abundance were measured by qPCR targeting: nitrifiers (AOA, ammonia oxidizing archaea; AOB, ammonia oxidizing bacteria), denitrifiers (*nirS*, cytochrome cd 1-containing nitrite reductase*; nosZI*, nitrous oxide reductase) and nitrogen fixers (*nifH*, nitrogenase reductase). All qPCR results are normalized to 16S rRNA copy numbers and presented as percent of the nucleic acid pool.

### Population and transcription dynamics for nitrogen related functional groups

Significant changes (ANOVA, p<0.05) in relative activity (mRNA abundance/16S rRNA gene abundance) were observed promptly between day 0 & 3 for all functional groups (except AOA and N-fixers in HM soil) in response to urea (Fig. 1). However, maximum relative transcription did not match maximum production peaks for corresponding substrates, or products, for each functional group. Nitrifiers (ammonia oxidizers) displayed niche differentiation, with time, length and strength of response differing between bacterial (AOB) and archaeal ammonia oxidizers (AOA). Relative activity of AOA increased (4.6-fold for LM and 1.6-fold for HM) under urea treatments at day 3 only, with a subsequent decrease (-19.3-fold for LM and -7-fold for HM) resulting in lower expression than in untreated soils (Fig. 1). AOB relative activity also increased but was sustained for a much longer period (3-63 days), with maximum activity (>11-fold change) seen at 21 and 35 days for HM+N and LM+N respectively (Fig. 1). Denitrifiers (both nitrite and nitrous oxide reducers) showed similar responses as AOA, with peak activity at day 3 and a rapid return to baseline, in the case of nitrite reducers decreasing to levels below those observed in non-urea treated soils (Fig. 1). To account for endogenous sources of N, N_2_ fixers were monitored through the activity of the nitrogenase gene (*nifH*). No significant changes were observed except for day 3 (LM +N only), with a subsequent decrease in activity below background. This decrease below background was observed for all N treated samples.

Changes in the relative contribution to total community composition were calculated by normalizing functional gene abundance to total 16S rRNA gene abundance per sample for each functional group (Fig. 1). The maximum observed relative abundance of each functional group differed for each group (HM and LM respectively): AOB, 19 & 12 %; AOA, 8 & 13 %; *nirS*, 6.3 & 2.9 %; *nosZI*, 3.3 & 3.4 %; *nifH*, 4.7 & 4.32 %. Further, large population changes over time were mostly limited to AOB. Generally, AOB comprised <1 % of the total community, but in response to urea increased up to 29-fold to make up 19 % (day 21 for HM) and 20-fold to make up 12 % (day 35 for LM) of the community in urea treated soils. In contrast, AOA were found at consistently high levels (median=4.2 %) in untreated soils, but numbers decreased >7-fold in response to urea (∼1.3 % at least 63 day). Similarly, other functional groups (*nosZI, nifH*) decreased or remained stable (*nirS*) in response to urea. Similar patterns for both activity and population changes were observed when absolute values were analyzed (Supplementary Fig. S2).

### N deposition induces both a genotypic and a transcriptional response at the community level that is modified by soil moisture content

Urea deposition imposed a general negative selective pressure leading to decreases in OTU level prokaryotic diversity (Shannon, -1.2-fold change), richness (- 1.5-fold change) and evenness (-1.1-fold change) at DNA level (Fig. 2a, Supplementary Fig. S3). The same pattern was observed when active microbes (based on RNA) were analyzed with decreases in OTU level prokaryotic diversity (Shannon, -1.3-fold change), richness (-1.9-fold change) and evenness (-1.2-fold change). Moisture was found to have a smaller, but significant, effect compared to urea, with LM samples consistently resulting in lower diversity and richness when compared to their HM pairs. Richness and diversity losses were not recovered even after 63 days. In contrast, samples where no urea was applied remained stable (i.e. constant diversity and richness).

**Fig. 2.**
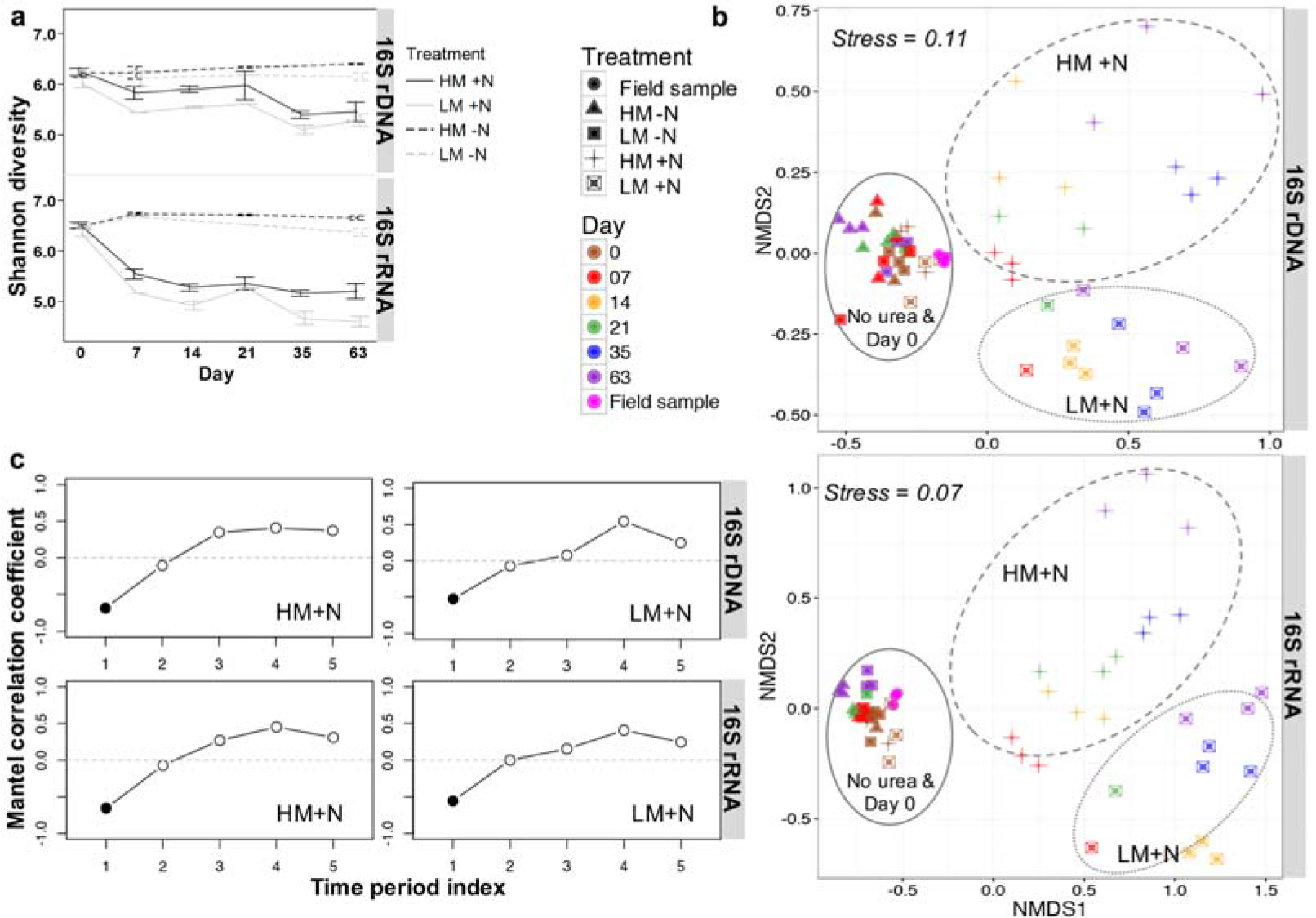
Total microbial community response (based on 16S rRNA gene amplicon profiling and clustering of sequences at OTU level (97% sequence similarity)) to urea (+/-1000 µg N/g dry soil) under two moisture conditions (LM = low moisture [- 10kPa]; HM = high moisture [-1.0kPa]) at both DNA and RNA level. Error bars are the standard error of the mean (n = 3, except day 7 [n=1; LM soil] and day 21 [n=1; LM soil]) for replicate mesocosms. (a) Changes in microbial diversity (Shannon) index over time in response to treatment. (b) Non-metric multidimensional scaling (NMDS) ordination plots based on Bray-Curtis distances showing relationships among samples based on OTU level changes in community composition. (c) Mantel correlogram showing autocorrelation on community composition by performing sequential Mantel tests between the Bray-Curtis dissimilarities and the grouping of samples using a time period index (index 1 represents 0-7 days; 2 represents 7-14; 3 represents 14-21; 4 represents 21-35; 5 represents 35-63). Filled circles represent significant correlation (p < 0.05) in community composition at specific time periods, with open circles indicating no significant correlation.

Urea deposition significantly altered community structure (Adonis test: F= 18.04, p< 0.001 for 16S rDNA and F= 26.27, p< 0.001 for 16S rRNA) as shown in a non-metric multidimensional scaling (NMDS) plot using a Bray-Curtis dissimilarity matrix (Fig. 2b and Supplementary Fig. S4). At both DNA and RNA level community changes along the first axis corresponded with changes in response to urea treatment, with the second axis accounting for changes in moisture. A pvclust analysis (hierarchical clustering with p-values calculated via multiscale bootstrap resampling, Supplementary Fig. S5) confirmed two major clusters [100% AU (Approximately Unbiased) and 100% BP (Bootstrap Probability)] formed by urea treated (HM+N and LM+N samples, excluding day 0), vs. untreated soils (HM-N, LM-N, field samples, and HM+N & LM+N at Day 0). Temporal variance within each cluster was confirmed using a Mantel correlogram analysis (Fig. 2c). Urea treated samples had significant changes in community composition immediately upon treatment (Day 0 to 7), with no return to baseline conditions by the end of the experiment. In contrast, untreated samples did not change significantly over time (Supplemental Fig. S6)

Changes in community structure were associated with shifts in major taxonomic lineages (Fig. 3). In general, phylum level changes in abundance and transcription where correlated to each other (Supplementary Table S2 and Fig. S7, S8). Urea deposition induced temporal changes in phylum level abundance with observed maximum fold changes per group (HM & LM at DNA level) being: Acidobacteria, -4.6 & -3.7; Actinobacteria, 2.4 & 5.3; Bacteroidetes, 4.6 & 2.2; Candidate Division WS3, -10.5 & -7; Chloroflexi, -2.9 & -2.6; Firmicutes, 10.8 & 16.2; Gemmatimonadetes, 2 & 3.3; Nitrospirae, -3.2 & -2; Planctomycetes, -3.7 & -2.5; Thaumarchaeota, -5.2 & -3.6; Verrucomicrobia, -2.5 & -2; Alphaproteobacteria, 1.4 & 1.7; Betaproteobacteria, 4 & 2; Deltaproteobacteria, -2.2 & -1.4; Gammaproteobacteria, 1.5 & 2.6. Normalized transcriptional activity (reads of 16S rRNA/reads of 16S rDNA) identified the Firmicutes and members within classes of the Proteobacteria as the most transcriptionally active. While abundant phyla tended to have high levels of normalized transcription, less abundant organisms like the Thaumarchaeota, were observed to have high normalized transcriptional activity especially under background conditions (Supplementary Fig. S7). In contrast, groups traditionally considered slow growers (e.g. Nitrospirae and Gemmatimonadetes) had low normalized transcription. It was also noted that while normalized transcription levels remained stable without urea, N deposition induced changes. These changes in normalized activity did not always match trends observed at individual DNA or RNA level (e.g. Firmicutes).

**Fig. 3.**
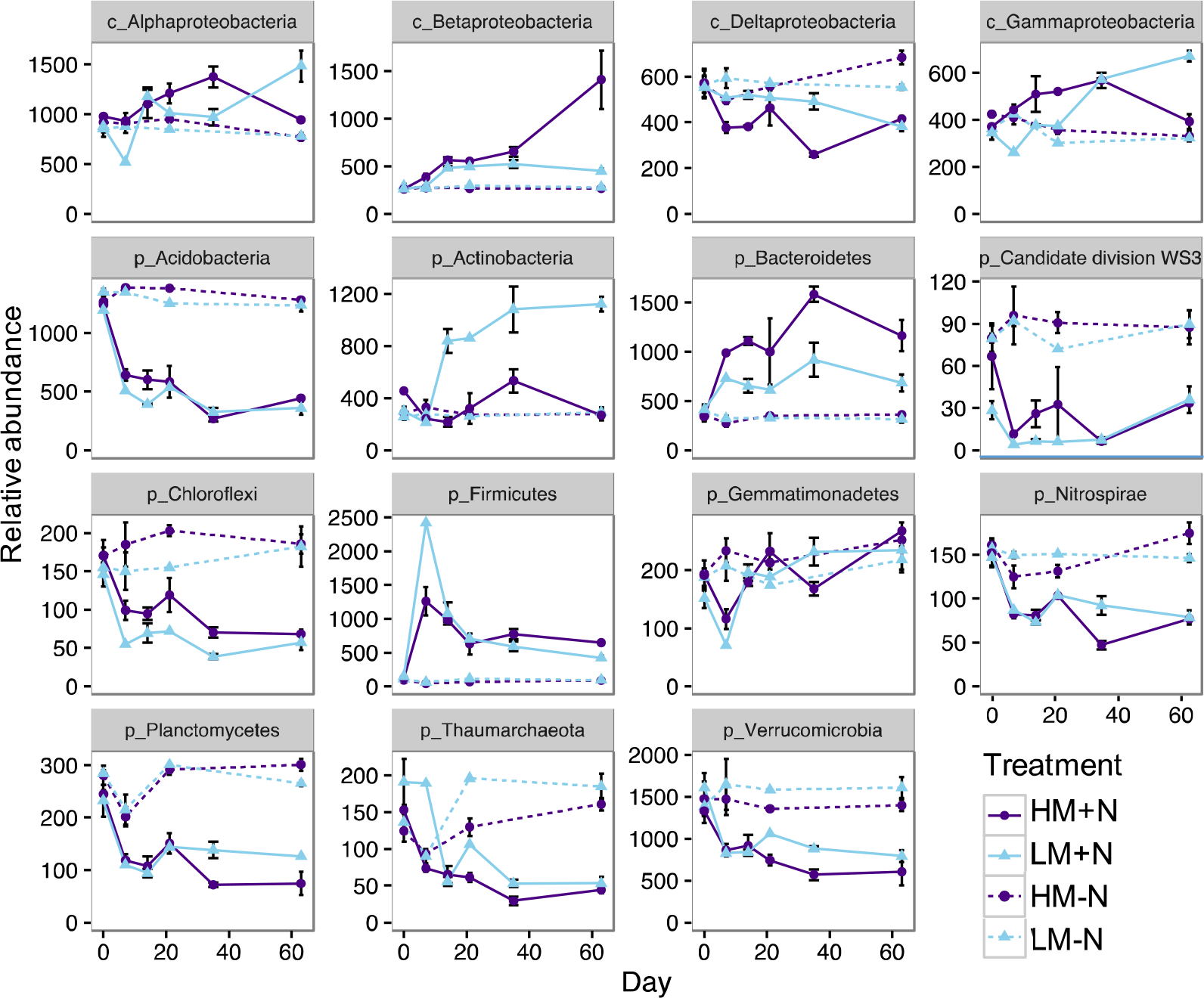
Phylum and class level (for Proteobacteria only) changes in abundance (DNA) representing relative contribution >1% of all detected phyla (based on OTUs clustered at 97% sequence similarity). A total of 7,400 sequences were examined per sample. Error bars are the standard error of the mean (n = 3, except day 7 [n=1; LM soil] and day 21 [n=1; LM soil]) for replicate mesocosms. Treatments = +/- N [+/-1000 µg N/g dry soil] under two moisture conditions (LM = low moisture [-10kPa]; HM = high moisture [-1.0kPa]). Abbreviations: c: Class; p: Phylum. See supplemental Fig. S8 for relative abundance

### Shifts in N and moisture status trigger OTU response linked to divergent life strategies

Since Fig. 3 only represents a taxonomic summary of all OTUs (irrespective of their response to treatments), it does not provide a clear indication of who is changing and why. To account for this, urea responsive OTUs were identified independently in RNA and DNA profiles (under each treatment) through a SIMPER analysis. OTUs accounting for 50% of the variance were analyzed (Fig. 4). Response patterns for detected OTUs were conserved between RNA and DNA profiles. However, while some OTUs responded similarly to urea under varying moisture conditions, marked differences were observed with no detectable pattern based on taxonomy.

**Fig. 4.**
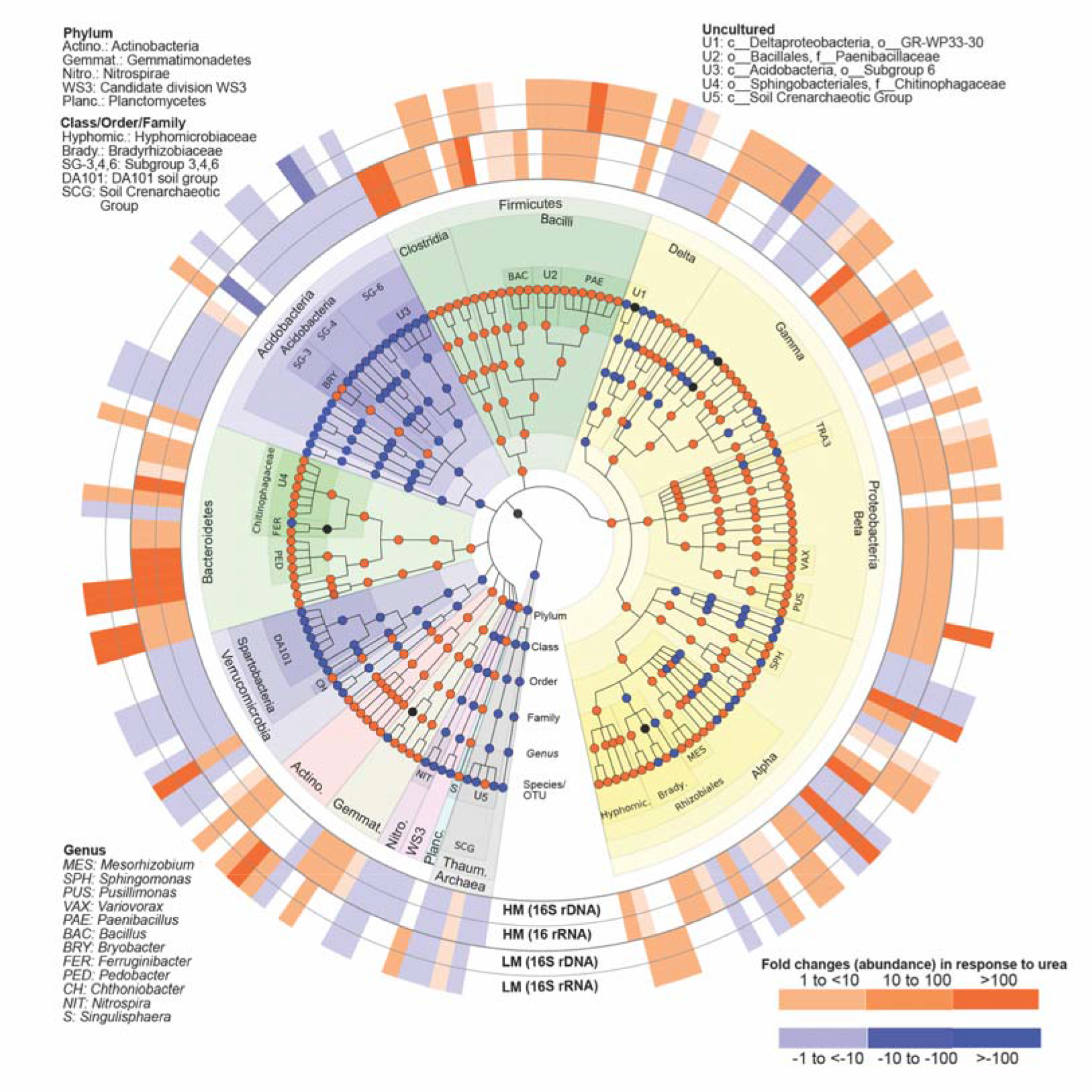
Taxonomic summary of OTUs responsive to urea treatment identified through similarity percentages (SIMPER) analysis (representing top 50% cumulative sum). The 4 outer rings represent fold changes in response to urea under high and low moisture content (MH & LM respectively) at either DNA or RNA level, with blank gaps indicating OTUs not identified in SIMPER analysis under the specified ring condition. Nodes on the tree (moving outwards from center) correspond to taxonomic level [Domain, Phylum, Class, Order, Family, Genus and Species/OTUs]. Nodes are colored based on dominant response (>50% conserved fold change response across OTUs within a node) with black notes indicating equal representation of positive and negatively responding OTUs. Shaded areas of branches delineate defined taxonomic groups. See Supplementary file (Table S3) for full classification.

OTUs within the Proteobacteria identified in the SIMPER analysis did not display a conserved response to urea, however when lower taxonomic levels were examined patterns emerged. A consistent positive response was seen for OTUs within the class Betaproteobacteria and the family Hyphomicrobiaceae, amongst others. Positive responses to urea where also observed at the phylum level for the Firmicutes, Bacteroidetes, Actinobacteria, Gemmatimonadetes and Planctomycetes, although the level of response varied across lower taxonomic levels. In contrast, with only some exceptions, OTUs within the phyla Acidobacteria, Verrucomicrobia, Nitrospirae, Candidate Division WS3 (also referred to as candidate phylum Latescibacteria) and the Thaumarchaeota all were negatively impacted by urea deposition.

To account for response patterns over time, we focused on OTUs that accounted for 30% of the variance in the SIMPER analysis (36 total), with individual OTU contributions ranging from 5 to 0.1 percent at the DNA level and 5 to 0.06 percent at the RNA level (Table S3). Temporal patterns were conserved between DNA and RNA profiles (Supplementary Fig. S9, S10), despite differences in absolute abundance. Once again, moisture acted as a modulator of response with the extent of impact dependent on the OTU (Fig. 5 and 6). While most functional groups responded immediately (at both DNA and RNA level), positively affected OTU responses were observed along all time points creating a succession of positively selected organisms. In contrast, negatively affected OTUs all responded within the first 2 time points indicating an immediate negative selective pressure (Fig. 6). Large variances in absolute changes were observed, even within similar organisms (e.g. Pedobacter), with fold changes ranging from -10.5 to 410 across both positively and negatively affected OTUs. Despite this, OTU response was noted to correspond to taxonomy, with both the effect (positive or negative) and the extent of response (fold change or total abundance) in line with predicted ecological growth strategies (r vs. k) predicted for different taxa. To test this, we predicted rRNA operon copy numbers (rrn) for all 36 OTUs and compared them to the observed maximum abundance, max fold change in population or observed growth rate per day. We consistently observed a non-linear response with an asymptote reached at higher copy numbers (Fig. 7). These trends were consistent independent of which moisture conditions were present at the time of response. To account for preferential response due to moisture, we selected the highest response for each organism and saw no clear difference in patterns. To account for potential biases due to uneven representation, OTUs were grouped into low (1-2 copies of rrn) or high (>2) copy number organisms (Supplementary Table S4). While significant changes (p < 0.05, Supplementary Fig. S11) were observed in most instances, exceptions were noted (e.g. growth rate under HM).

**Fig. 5.**
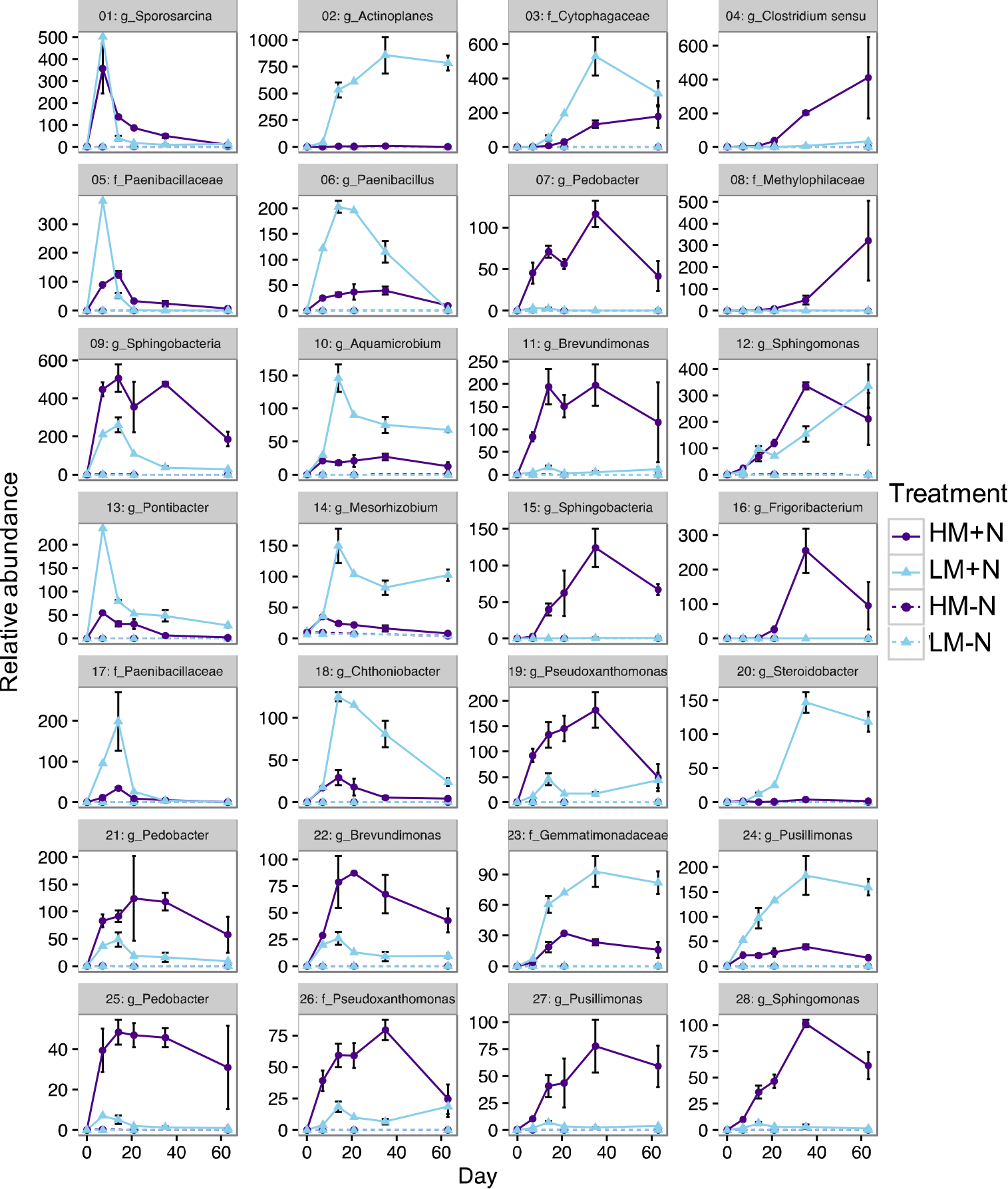
Population (16S rDNA) changes (abundance based on 7400 reads per samples) for OTUs identified as positively responsive to urea treatment based on similarity percentages (SIMPER) analysis (representing top 30% cumulative sum). Treatments = +/- N [+/-1000 µg N/g dry soil] under two moisture conditions (LM = low moisture [-10kPa]; HM = high moisture [-1.0kPa]).

**Fig. 6.**
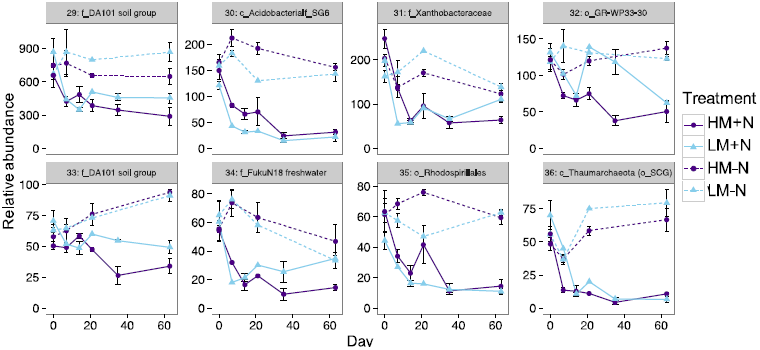
Population (16S rDNA) changes (abundance based on 7,400 reads per samples) for OTUs identified as negatively responsive to urea treatment based on similarity percentages (SIMPER) analysis (representing top 30% cumulative sum). Treatments = +/- N [+/-1000 µg N/g dry soil] under two moisture conditions (LM = low moisture [-10kPa]; HM = high moisture [-1.0kPa]).

**Fig. 7.**
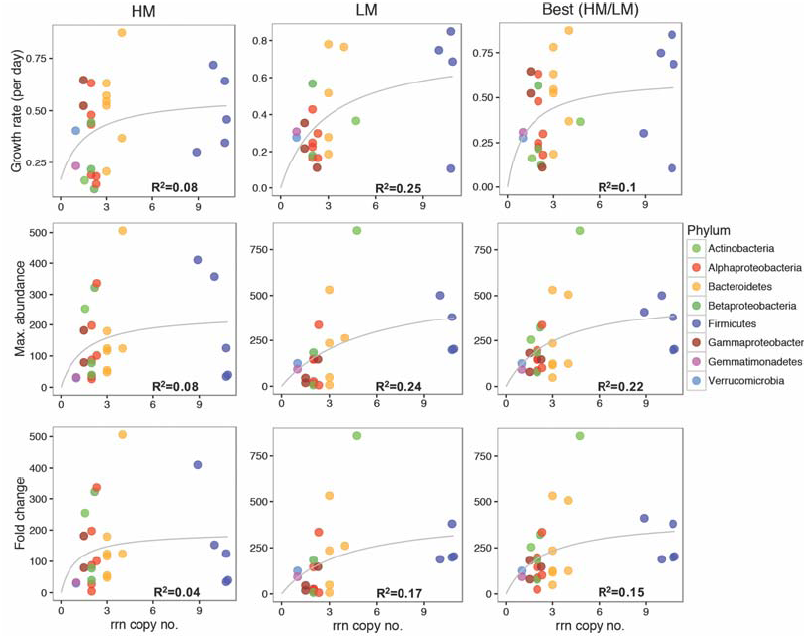
Relationship between predicted ribosomal RNA operon (rrn) copy numbers and growth rate (per day), maximum observed population change, or fold change in response to N treatment under both high moisture (HM) content, low moisture (LM) content and best growth either in HM or in LM (based on maximum observed growth). Copy number was estimated using rrn database (Stoddard *et al*. 2015). Copy number values were obtained by finding the closest match (lowest taxonomic level possible) to each OTU and retrieving the mean rRNA copy number for that group.

## Discussion

Functional profiling (identification and quantification of specific functional genes/transcripts) is normally utilized to link chemical transformations to specific microbial populations capable of catalyzing reactions. However, functional groups are comprised of taxonomically diverse species of microbes with different lifestyle strategies that are unlikely to share a conserved response to an ecosystem disturbance (Ho *et al*. 2012). While functional profiling allows us to measure the net response of a functional group, and could serve as a proxy for determining the importance of the group in a sample, it does not identify how specific organisms benefit from a catalyzed transformation. Here we used a controlled microcosm experiment to measure the response of soil communities to a disturbance in the form of changes in moisture and nitrogen (urea) deposition. Functional analysis (qPCR) demonstrated a biological response to urea, but differing responses to moisture depending on group (Fig. 1). Responses are potentially linked to different life strategies amongst these groups. Ammonia oxidizers displayed contrasting population and expression profiles, suggesting niche differentiation driven by time and/or substrate concentration. AOA responded early, and declined as new N was made available while AOB responded later with population swings spanning from near detection limit to most dominant group. These observations match prior reports showing AOA prefer low N concentrations, while AOB respond vigorously to N deposition (Di *et al*. 2010; Sterngren *et al*. 2015). This has been interpreted as evidence for differing lifestyles for AOB and AOA, with AOA preferring nutrient poor conditions and AOB dominating in rich ones (Sterngren *et al*. 2015). However, prior assertions that AOB are solely important for driving nitrification might be overstated given that transcriptional activity for both groups is comparable if compared at peak time (Di *et al*. 2009). This contrasting use of energy between functionally redundant organisms might explain the low correlations between processes and the abundance of their respective functional populations (Rocca *et al*. 2015). When we examine the response of other functional groups benefiting from influxes of N, like denitrifiers, we see no significant change in population sizes suggesting that either energy is being utilized for physiological maintenance or otherwise for redox balance/homeostasis (Hartsock & Shapleigh 2011; Li *et al*. 2012; Dietrich *et al*. 2013). The distinction here being that we use the term physiological maintenance as it refers to the state of energetics in a cell where the energy consumed is used for functions other than the production of new cell material (i.e. growth) (van Bodegom 2007; Lipson 2015). Alternatively, redox balance reactions are used to maintain viable metabolic processes by controlling the redox state of all the cellular components (Green & Paget 2004). In contrast, organism adapted to low N concentrations, like N fixers, decline in response to exogenous N demonstrating real time selective pressure in a complex ecosystem. These responses also highlight the temporal nature of these relationships and how by following niche differentiation high number of functionally redundant organisms can be maintained (Stempfhuber *et al*. 2016). However, the use of very high concentrations of urea (leading to rapid hydrolysis to ammonium followed by substantial nitrification) has major consequences for soil pH, physicochemical parameters, and potentially other factors (e.g. osmolarity). Without accounting for those it is unclear what the direct mechanism causing an increase or decrease in the relative abundance of a specific population is.

Despite this, our observations highlight how lifestyle preferences for organisms may be reflected in their dominance in the ecosystem. Prior work suggests that AOA dominate in soils with low N inputs, but AOB numbers are higher at times of high N loading or in ecosystems with consistent N deposition (Gong *et al*. 2013; Venterea *et al*. 2015; Sterngren *et al*. 2015; Li *et al*. 2016). This would suggest that a dynamic ecosystem with varying nutrient levels would select for a higher diversity of organisms that maintain ecosystem processes stable over time and space (Wang and Loreau, 2014). Indeed, our data supports this with alpha diversity (calculated based on 16S amplicon analysis at both DNA and RNA) decreasing in response to urea. This is inconsistent with plant responses to nutrient deposition in which multiple resources need to be added to elicit a response (Harpole *et al*. 2016), although contrasting results have been observed (Suding *et al*. 2005; Bai *et al*. 2010; Song *et al*. 2011; 2012). For microbes, high site to site variance is reported (De Schrijver *et al*. 2011; Leff *et al*. 2015), but similar negative responses are suggested and could be linked to increased competition in the absence of natural ecosystem variability. However, links between microbial and plant response suggest interplay between the response of macro and microbiota (Zeng *et al*. 2016). While previous work suggests an important role for moisture in controlling community composition (Waldrop & Firestone 2006), we only observed a modifier role in our experiment.

Although broad observations align with ecological theory, precise identification of responsive organisms is rarely carried out. Here we note that while at phylum level clear responses (+/- fold change) are observed, variance is seen at the OTU level suggesting intra-taxonomic (i.e. same phylum but different species or OTUs) diversity. We hypothesized this reflects the life history strategies of the different organisms. Attempts to link specific transformations to organisms failed, potentially due to the succession of functionally redundant organisms that respond at different time with non-overlapping optima. That is, while functional gene abundance provides the population size of organisms capable of carrying out a process, the group may be composed of many OTUs with divergent life strategies or metabolic potentials that affect when they can respond. This makes functional gene measurement an average of all OTU subpopulations carrying that gene. However, community response allows us to identify OTUs responsive to N deposition, which when analyzed independently, provides insights into metabolic preferences (i.e. aerobic vs. anaerobic, nitrifier vs. denitrifier) based on time and response to treatments. Taxonomic groups regularly recognized as native to, or abundant in, oligotrophic conditions declined in the presence of urea. Most of these groups are still poorly understood, and included the Acidobacteria, Verrucomicrobia, Nitrospirae, Candidate Division WS3 (also referred to as candidate phylum Latescibacteria) and the Thaumarchaeota. These organisms are predicted to be slow growers with the Thaumarchaeal response confirming the AOA patterns observed at the functional level. In contrast, positively responding organisms are those generally associated with groups considered eutrophic or capable of fast response. This discrepancy based on life history strategies has been proposed and applied to microbes previously, and suggests that an organisms’ ability to grow, utilize carbon, generate proteins and efficiently transform resources to biomass, amongst others, is related to its rRNA operon copy number (Klappenbach *et al*. 2000; Stevenson & Schmidt 2004; Dethlefsen & Schmidt 2007; Roller *et al*. 2016). When applied to communities, it is associated with microbial successions in which decreases in copy numbers are associated with later stages of succession including in soils (Nemergut *et al*. 2015). For example, two OTUs matching the Verrucomicrobial OTU DA101 where found to be negatively affected by urea, and at least one was found to be highly abundant under background conditions. DA101 seems to be a common soil (and grassland) organism identified throughout the world (Felske & Akkermans 1998; O’Farrell & Janssen 1999; Brewer *et al*. 2016). Based on growth (Sangwan *et al*. 2005) and genome reconstructions (Brewer *et al*. 2016), these organisms are predicted to be slow but efficient growers (k strategists). In contrast, most of the positively affected organisms seemed to posses higher *rrn* copy numbers and included members of the Proteobacteria and Bacteroidetes in line with prior predictions (Fierer *et al*. 2007). Statistical analysis supported this interpretation with low copy numbers (1-2) significantly associated to a negative response to N deposition, while high copy numbers (>2) were linked to increased capacity for growth, growth rate and maximum abundance. However, we found a non-linear relationship between increased *rrn* copy numbers and growth capacity, best fitted by models reaching an asymptote. These are first order models that suggest that while a benefit exists where increased copy numbers lead to increased growth rate, after a certain threshold other variables might limit any benefit. Alternatively, a decrease in growth rate might be observed with increasing copy numbers once a tradeoff threshold is passed (Lipson 2015). However, when *rrn* copy numbers are log2 transformed, a significant linear fit was observed as seen in prior studies (Roller *et al*. 2016). In our study these predictions are made complicated due to the observed intra-taxonomic variance that can arise from the lack of accurate knowledge of copy numbers for many organisms, or from metabolic plasticity at higher taxonomic levels. In addition, our analysis focused on N responsive organisms only, and with only 38 identified it indicates that most organisms were neither positively nor negatively affected. This could explain why certain organisms (e.g. Actinobacteria) expected to be k strategist, based on their ability to produce secondary metabolites (Abdelmohsen *et al*. 2015) and compete with other organisms (Barka *et al*. 2015), showed a positive response to N deposition. Alternatively, the low number of responsive organisms could indicate that our false discovery rate corrections were too restrictive.

These findings help us get closer to understanding not just the metabolic potential of organisms in soils, but the role specific pathways play for an organism. It also allows us to understand the repercussion of disturbances and management of soils on below ground biodiversity. The knowledge gained through these type of observations, and integration of life history strategies into microbial ecology, will get us one step closer to microbiome management as part of soil care.

## Acknowledgements

This work was funded by the New Zealand Government through the New Zealand Fund for Global Partnerships in Livestock Emissions Research to support the objectives of the Livestock Research Group of the Global Research Alliance on Agricultural Greenhouse Gases (Agreement number: 16084) awarded to SEM and the University of Otago.

## Supporting information

**Fig. S1** N_2_O response in soils treated with urea (+/- 1000 µg N/g dry soil) under two moisture conditions (LM = low moisture [-10kPa]; HM = high moisture [-1.0kPa]). Error bars are the standard error of the mean (n ≥ 3) for replicate mesocosms.

**Fig. S2** Functional group response (absolute quantification) in soils treated with urea (+/-1000 µg N/g dry soil) under two moisture conditions (LM = low moisture [- 10kPa]; HM = high moisture [-1.0kPa]). Gene and transcript abundance were measured from DNA template (1 ng of DNA) and cDNA template (1 ng RNA). Error bars are the standard error of the mean (n = 3, except day 7 [n=1; LM soil] and day 21 [n=1; LM soil]) for replicate mesocosms. Absolute gene and transcript abundance were measured by qPCR targeting: 16S (total prokaryotic community), nitrifiers (AOA, ammonia oxidizing archaea; AOB, ammonia oxidizing bacteria), denitrifiers (*nirS*, cytochrome cd1-containing nitrite reductase*; nosZI*, nitrous oxide reductase) and nitrogen fixers (*nifH*, nitrogenase reductase).

**Fig. S3** Changes in microbial a) Richness and b) Evenness (Pielou’s) over time in response to treatment.

**Fig. S4** Stress plots for Fig. 2b.

**Fig. S5** Pvclust tree displaying sample clustering based on Bray-Curtis distances calculated from 16S rRNA gene community composition and indicating significant clusters based on p values ([AU (approximately unbiased) BP (bootstrap probability)]) for each node. Red boxes mark clusters with 95% confidence. Bootstrap replication (n=1000). Two clusters: with urea (light red box) and no urea + day 0 N treated samples (light green box).

**Fig. S6** Mantel correlogram showing autocorrelation on community composition by performing sequential Mantel tests between the Bray-Curtis dissimilarities and the grouping of samples using a time period index (index 1 represents 0-7 days; 2 represents 7-21; 3 represents 21-63). Opened circles represent no significant correlations (p > 0.05) in community composition at specific time periods.

**Fig. S7** Changes in abundance (DNA), activity (RNA) and RNA/DNA ratio for phyla, or classes, representing top 11 phyla (based on OTUs clustered at 97% sequence similarity). A total of 7,400 sequences were examined per sample. Error bars are the standard error of the mean (n = 3, except day 7 [n=1; LM soil] and day 21 [n=1; LM soil]) for replicate mesocosms. Treatments = +/- N [+/- 1000 µg N/g dry soil] under two moisture conditions (LM = low moisture [-10kPa]; HM = high moisture [- 1.0kPa]). Abbreviations: Firmi., Firmicutes; Verru., Verrucomicrobia; Bact., Bacteroidetes; Acido., Acidobacteria; Actino., Actinobacteria; Planct., Planctomycetes; Gemma., Gemmatimonadetes; Thaum., Thaumarchaeota; Chloro., Chloroflexi, Nitro., Nitrospirae.

**Fig. S8** Phylum level changes (relative abundance) in genome (16S rDNA) and transcript (16S rRNA) levels representing relative contribution >1% of all detected phyla (based on OTUs clustered at 97% sequence similarity). A total of 7,400 sequences were examined per sample. Treatments = +/- N [+/- 1000 µg N /g dry soil] under two moisture conditions (LM = low moisture [-10kPa]; HM = high moisture [- 1.0kPa]).

**Fig. S9** Transcriptional (16S rRNA) and population (16S rDNA) changes (absolute abundance based on 7400 reads per samples) for OTUs identified as positively responsive to urea treatment based on similarity percentage (SIMPER) analysis (representing top 30% cumulative sum). Treatments = +/- N [+/- 1000 µg N/g dry soil] under two moisture conditions (LM = low moisture [-10kPa]; HM = high moisture [-1.0kPa]).

**Fig. S10** Transcriptional (16S rRNA) and population (16S rDNA) changes (absolute abundance based on 7400 reads per samples) for OTUs identified as negatively responsive to urea treatment based on similarity percentage (SIMPER) analysis (representing top 30% cumulative sum). Treatments = +/- N [+/- 1000 µg N (urea)/g dry soil] under two moisture conditions (LM = low moisture [-10kPa]; HM = high moisture [-1.0kPa]).

**Fig. S11** Relationship between predicted ribosomal RNA operon (rrn) copy numbers and observed growth rate (per day), maximum observed population change, or fold change in population abundance for OTUs responsive to N treatment under both high moisture (HM) content. Copy number was estimated using rrn database (Stoddard *et al*. 2015). Predicted rrn copy numbers represent the mean rRNA copy number for the closest taxonomic match (at the lowest taxonomic level possible) for each OTU. The rrn copy numbers were log2 transformed before linear regression analysis. Significant “p” value is marked with an asterisk (*p<0.05; **p<0.01; ***p<0.001)

**Table S1** Primer pairs used in this study

**Table S2** Pairwise correlation between observed phylum (or class) abundance at DNA and RNA level for urea (+N) treated soils. Correlation analysis was done between DNA (16S rDNA) and RNA (16S rRNA) samples based on mean absolute abundance (per 7,400 sequence reads) at each time point (day 0, 7, 14, 21, 35, 63). Only Proteobacteria shown at class level.

**Table S3** Top OTUs cumulatively contributing 50% of the variance between groups (+Urea; −Urea) at 16S rDNA and 16S rRNA levels based on SIMPER analsysis.

**Table S4:** Two sample t-test for mean comparison between low copy number rrn (rRNA operon) samples (1-2) and high copy number of rrn samples (>2). The significant correlation (p<0.05) are showed as bold.

## References

Abdelmohsen UR, Grkovic T, Balasubramanian S et al. (2015) Elicitation of secondary metabolism in actinomycetes. Biotechnology Advances, 33, 798–811.

Altschul SF, Gish W, Miller W, Myers EW, Lipman DJ (1990) Basic local alignment search tool. Journal of Molecular Biology, 215, 403–410.

Avrahami S, Liesack W, Conrad R (2003) Effects of temperature and fertilizer on activity and community structure of soil ammonia oxidizers. Environmental Microbiology, 5, 691–705.

Bai Y, Wu J, Clark CM, Naeem S, Pan Q (2010) Tradeoffs and thresholds in the effects of nitrogen addition on biodiversity and ecosystem functioning: evidence from inner Mongolia Grasslands. Global Change Biology, 16, 358–372.

Baral KR, Thomsen AG, Olesen JE, Petersen SO (2014) Controls of nitrous oxide emission after simulated cattle urine deposition. Agriculture, Ecosystems & Environment, 188, 103–110.

Barka EA, Vatsa P, Sanchez L et al. (2015) Taxonomy, Physiology, and Natural Products of Actinobacteria. Microbiology and Molecular Biology Reviews, 80, 1– 43.

Bayer B, Vojvoda J, Offre P et al. (2015) Physiological and genomic characterization of two novel marine thaumarchaeal strains indicates niche differentiation. 10, 1051–1063.

Brenzinger K, Dörsch P, Braker G (2015) pH-driven shifts in overall and transcriptionally active denitrifiers control gaseous product stoichiometry in growth experiments with extracted bacteria from soil. Frontiers in microbiology, 6, 1226.

Brewer T, Handley K, Carini P, Gibert J, Fierer N (2016) Genome reduction in an abundant and ubiquitous soil bacterial lineage.

Butterbach-Bahl K, Baggs EM, Dannenmann M, Kiese R, Zechmeister-Boltenstern S (2013) Nitrous oxide emissions from soils: how well do we understand the processes and their controls? Philosophical Transactions of the Royal Society B: Biological Sciences, 368, 20130122–20130122.

Caporaso JG, Kuczynski J, Stombaugh J et al. (2010) QIIME allows analysis of high-throughput community sequencing data. Nature methods, 7, 335–336.

Caporaso JG, Lauber CL, Walters WA et al. (2012) Ultra-high-throughput microbial community analysis on the Illumina HiSeq and MiSeq platforms. The ISME Journal, 6, 1621–1624.

Clough, TJ, Lanigan, GJ, de Klein, CAM et al. (n.d.). Influence of soil moisture on co-denitrification fluxes from a urea-affected pasture soil. In Review.

de Klein CAM, Luo J, Woodward KB et al. (2014a) The effect of nitrogen concentration in synthetic cattle urine on nitrous oxide emissions. Agriculture, Ecosystems & Environment, 188, 85–92.

de Klein CA, Shepherd MA, van der Weerden TJ (2014b) Nitrous oxide emissions from grazed grasslands: interactions between the N cycle and climate change — a New Zealand case study. Current Opinion in Environmental Sustainability, 9-10, 131–139.

De Schrijver A, De Frenne P, Ampoorter E et al. (2011) Cumulative nitrogen input drives species loss in terrestrial ecosystems. Global Ecology and Biogeography, 20, 803–816.

Delmont TO, Eren AM, Maccario L et al. (2015) Reconstructing rare soil microbial genomes using in situ enrichments and metagenomics. Frontiers in microbiology, 6, 1–15.

Dethlefsen L, Schmidt TM (2007) Performance of the translational apparatus varies with the ecological strategies of bacteria. Journal of bacteriology, 189, 3237– 3245.

Di HJ, Cameron KC, Shen JP et al. (2009) Nitrification driven by bacteria and not archaea in nitrogen-rich grassland soils. Nature Geoscience, 2, 621–624.

Di HJ, Cameron KC, Shen J-P et al. (2010) Ammonia-oxidizing bacteria and archaea grow under contrasting soil nitrogen conditions. FEMS Microbiology Ecology, 72, 386–394.

Dietrich LEP, Okegbe C, Price-Whelan A et al. (2013) Bacterial Community Morphogenesis Is Intimately Linked to the Intracellular Redox State. Journal of bacteriology, 195, 1371–1380.

Edgar RC (2010) Search and clustering orders of magnitude faster than BLAST. Bioinformatics, 26, 2460–2461.

Felske, Akkermans (1998) Prominent occurrence of ribosomes from an uncultured bacterium of the Verrucomicrobiales cluster in grassland soils. Letters in applied microbiology, 26, 219–223.

Fierer N, Bradford MA, Jackson RB (2007) Toward an ecological classification of soil bacteria. Ecology, 88, 1354–1364.

Gardiner MJ, Radford T (1980) Soil associations of Ireland and their land use potential: explanatory bulletin to soil map of Ireland 1980. An Foras Taluntais Dublin.

Gong P, Zhang L, Wu Z, Shang Z, Li D (2013) Does the nitri cation inhibitor dicyandiamide affect the abundance of ammonia-oxidizing bacteria and archaea in a Hap-Udic Luvisol? Journal of Soil Science and Plant Nutrition, 13, 35–42.

Green J, Paget MS (2004) Bacterial redox sensors. Nature reviews microbiology, 2, 954–966.

Gubry-Rangin C, Kratsch C, Williams TA et al. (2015) Coupling of diversification and pH adaptation during the evolution of terrestrial Thaumarchaeota. Proceedings of the National Academy of Sciences, 112, 9370–9375.

Hamonts K, Balaine N, Moltchanova E et al. (2013) Influence of soil bulk density and matric potential on microbial dynamics, inorganic N transformations, N_2_O and N_2_ fluxes following urea deposition. Soil Biology & Biochemistry, 65, 1–11.

Harpole WS, Sullivan LL, Lind EM et al. (2016) Addition of multiple limiting resources reduces grassland diversity. Nature, 537, 93–96.

Hartman AL, Lough DM, Barupal DK et al. (2009) Human gut microbiome adopts an alternative state following small bowel transplantation. Proceedings of the National Academy of Sciences, 106, 17187–17192.

Hartsock A, Shapleigh JP (2011) Physiological Roles for Two Periplasmic Nitrate Reductases in Rhodobacter sphaeroides 2.4.3 (ATCC 17025). Journal of bacteriology, 193, 6483–6489.

Hatzenpichler R (2012) Diversity, physiology, and niche differentiation of ammonia-oxidizing archaea. Applied and Environmental Microbiology, 78, 7501–7510.

Henry S, Bru D, Stres B, Hallet S, Philippot L (2006) Quantitative detection of the *nosZ* gene, encoding nitrous oxide reductase, and comparison of the abundances of 16S rRNA, *narG, nirK*, and nosZ genes in soils. Applied and Environmental Microbiology, 72, 5181–5189.

Herrero M, Henderson B, Havlík P et al. (2016) Greenhouse gas mitigation potentials in the livestock sector. Nature Climate Change, 6, 452–461.

Ho A, Kerckhof F-M, Luke C et al. (2012) Conceptualizing functional traits and ecological characteristics of methane-oxidizing bacteria as life strategies. Environmental Microbiology Reports, 5, 335–345.

Jia Z, Conrad R (2009) Bacteriarather than Archaea dominate microbial ammonia oxidation in an agricultural soil. Environmental Microbiology, 11, 1658–1671.

Klappenbach JA, Dunbar JM, Schmidt TM (2000) rRNA Operon Copy Number Reflects Ecological Strategies of Bacteria. Applied and Environmental Microbiology, 66, 1328–1333.

Leff JW, Jones SE, Prober SM et al. (2015) Consistent responses of soil microbial communities to elevated nutrient inputs in grasslands across the globe. Proceedings of the National Academy of Sciences, 112, 10967–10972.

Leininger S, Urich T, Schloter M et al. (2006) Archaea predominate among ammonia-oxidizing prokaryotes in soils. Nature, 442, 806–809.

Li C, Di HJ, Cameron KC, Podolyan A, Zhu B (2016) Effect of different land use and land use change on ammonia oxidiser abundance and N_2_O emissions. Soil Biology & Biochemistry, 96, 169–175.

Li Y, Katzmann E, Borg S, Schuler D (2012) The Periplasmic Nitrate Reductase Nap Is Required for Anaerobic Growth and Involved in Redox Control of Magnetite Biomineralization in Magnetospirillum gryphiswaldense. Journal of bacteriology, 194, 4847–4856.

Lipson DA (2015) The complex relationship between microbial growth rate and yield and its implications for ecosystem processes. Frontiers in microbiology, 6, 1–5.

Liu B, Liu B, Frostegard Å et al. (2014) Impaired Reduction of N_2_O to N2 in Acid Soils Is Due to a Posttranscriptional Interference with the Expression of nosZ. mBio, 5, e01383–14–e01383–14.

Liu B, Mørkved PT, Frostegård Å, Bakken LR (2010) Denitrification gene pools, transcription and kinetics of NO, N_2_O and N2 production as affected by soil pH. FEMS Microbiology Ecology, 72, 407–417.

McMurdie PJ, Holmes S (2013) phyloseq: an R package for reproducible interactive analysis and graphics of microbiome census data. PLoS ONE.

Morales SE, Holben WE (2013) Functional response of a near-surface soil microbial community to a simulated underground CO_2_ storage leak (MR Mormile, Ed,). PLoS ONE, 8, e81742–10.

Nemergut DR, Knelman JE, Ferrenberg S et al. (2015) Decreases in average bacterial community rRNA operon copy number during succession. 10, 1147–1156.

Nicol GW, Leininger S, Schleper C, Prosser JI (2008) The influence of soil pH on the diversity, abundance and transcriptional activity of ammonia oxidizing archaea and bacteria. Environmental Microbiology, 10, 2966–2978.

Oksanen J, Blanchet FG, Kindt R, Legendre P Vegan: Community Ecology Package (2013) R package version 2.0-7. https://cran.r-project.org/package=vegan.

O’Farrell KA, Janssen PH (1999) Detection of Verrucomicrobia in a Pasture Soil by PCR-Mediated Amplification of 16S rRNA Genes. Applied and Environmental Microbiology, 65, 4280–4284.

Pratscher J, Dumont MG, Conrad R (2011) Ammonia oxidation coupled to CO_2_ fixation by archaea and bacteria in an agricultural soil. In:, pp. 4170–4175.

Prosser JI, Nicol GW (2012) Archaeal and bacterial ammonia-oxidisers in soil: the quest for niche specialisation and differentiation. Trends in Microbiology, 20, 523–531.

Quast C, Pruesse E, Yilmaz P et al. (2012) The SILVA ribosomal RNA gene database project: improved data processing and web-based tools. Nucleic Acids Research, 41, D590–D596.

R Development Core Team (2008) R: A language and environment for statistical computing. R Foundation for Statistical Computing, Vienna, Austria.

Rinke C, Schwientek P, Sczyrba A et al. (2013) Insights into the phylogeny and coding potential of microbial dark matter. Nature, 499, 431–437.

Rocca JD, Hall EK, Lennon JT et al. (2015) Relationships between protein-encoding gene abundance and corresponding process are commonly assumed yet rarely observed. 9, 1693–1699.

Roller BRK, Stoddard SF, Schmidt TM (2016) Exploiting rRNA operon copy number to investigate bacterial reproductive strategies. Nature Microbiology, 1–7.

Rotthauwe JH, Witzel KP, Liesack W (1997) The ammonia monooxygenase structural gene *amoA* as a functional marker: molecular fine-scale analysis of natural ammonia-oxidizing populations. Applied and Environmental Microbiology, 63, 4704–4712.

Rösch C, Bothe H (2005) Improved assessment of denitrifying, N_2_-fixing, and total-community bacteria by terminal restriction fragment length polymorphism analysis using multiple restriction enzymes. Applied and Environmental Microbiology, 71, 2026–2035.

Saggar S, Jha N, Deslippe J et al. (2013) Denitrification and N_2_O:N2 production in temperate grasslands: Processes, measurements, modelling and mitigating negative impacts. Science of The Total Environment, 465, 173–195.

Sangwan P, Kovac S, Davis KER, Sait M, Janssen PH (2005) Detection and cultivation of soil verrucomicrobia. Applied and Environmental Microbiology, 71, 8402–8410.

Song L, Bao X, Liu X, Zhang F (2012) Impact of nitrogen addition on plant community in a semi-arid temperate steppe in China. Journal of Arid Land, 4, 3– 10.

Song L, Bao X, Liu X et al. (2011) Nitrogen enrichment enhances the dominance of grasses over forbs in a temperate steppe ecosystem. Biogeosciences, 8, 2341– 2350.

Spiro S (2012) Nitrous oxide production and consumption: regulation of gene expression by gas-sensitive transcription factors. Philosophical Transactions of the Royal Society B: Biological Sciences, 367, 1213–1225.

Stahl DA, la Torre de JR (2012) Physiology and diversity of ammonia-oxidizing archaea. Annual Review of Microbiology, 66, 83–101.

Stempfhuber B, Richter-Heitmann T, Regan KM et al. (2016) Spatial Interaction of Archaeal Ammonia-Oxidizers and Nitrite-Oxidizing Bacteria in an Unfertilized Grassland Soil. Frontiers in microbiology, 6, 861–15.

Sterngren AE, Hallin S, Bengtson P (2015) Archaeal ammonia oxidizers dominate in numbers, but bacteria drive gross nitrification in N-amended grassland soil. Frontiers in microbiology, 6, 1620–8.

Stevenson BS, Schmidt TM (2004) Life History Implications of rRNA Gene Copy Number in Escherichia coli. Applied and Environmental Microbiology, 70, 6670– 6677.

Stoddard SF, Smith BJ, Hein R, Roller BRK, Schmidt TM (2015) rrnDB: improved tools for interpreting rRNA gene abundance in bacteria and archaea and a new foundation for future development. Nucleic Acids Research, 43, D593–D598.

Suding KN, Collins SL, Gough L et al. (2005) Functional- and abundance-based mechanisms explain diversity loss due to N fertilization. Proceedings of the National Academy of Sciences, 102, 4387–4392.

Suzuki R, Shimodaira H (2006) Pvclust: an R package for assessing the uncertainty in hierarchical clustering. Bioinformatics, 22, 1540–1542.

Taylor AE, Zeglin LH, Wanzek TA, Myrold DD, Bottomley PJ (2012) Dynamics of ammonia-oxidizing archaea and bacteria populations and contributions to soil nitrification potentials. 6, 2024–2032.

Throbäck IN, Enwall K, Jarvis Å, Hallin S (2004) Reassessing PCR primers targeting nirS, *nirK* and *nosZ* genes for community surveys of denitrifying bacteria with DGGE. FEMS Microbiology Ecology, 49, 401–417.

Tourna M, Freitag TE, Nicol GW, Prosser JI (2008) Growth, activity and temperature responses of ammonia-oxidizing archaea and bacteria in soil microcosms. Environmental Microbiology, 10, 1357–1364.

van Bodegom P (2007) Microbial Maintenance: A Critical Review on Its Quantification. Microbial Ecology, 53, 513–523.

Venterea RT, Clough TJ, Coulter JA et al. (2015) Ammonium sorption and ammonia inhibition of nitrite-oxidizing bacteria explain contrasting soil N_2_O production. Scientific Reports, 5, 12153–15.

Waldrop MP, Firestone MK (2006) Response of microbial community composition and function to soil climate change. Microbial Ecology, 52, 716–724.

Wei W, Isobe K, Nishizawa T et al. (2015) Higher diversity and abundance of denitrifying microorganisms in environments than considered previously. 1–12.

Yergeau E, Kang S, He Z, Zhou J, Kowalchuk GA (2007) Functional microarray analysis of nitrogen and carbon cycling genes across an Antarctic latitudinal transect. 1, 163–179.

Zeng J, Liu X, Song L et al. (2016) Nitrogen fertilization directly affects soil bacterial diversity and indirectly affects bacterial community composition. Soil Biology & Biochemistry, 92, 41–49.

